# Tandem duplications lead to loss of fitness effects in CRISPR-Cas9 data

**DOI:** 10.1101/325076

**Authors:** Emanuel Gonçalves, Fiona M Behan, Sandra Louzada, Damien Arnol, Euan Stronach, Fengtang Yang, Kosuke Yusa, Oliver Stegle, Francesco Iorio, Mathew J Garnett

## Abstract

CRISPR-Cas9 gene-editing is widely used to study gene function and is being advanced for therapeutic applications. Structural rearrangements are a ubiquitous feature of cancers and their impact on CRISPR-Cas9 gene-editing has not yet been systematically assessed. Utilising CRISPR-Cas9 knockout screens for 163 cancer cell lines, we demonstrate that targeting tandem amplified regions is highly detrimental to cellular fitness, in contrast to amplifications caused by chromosomal duplications which have little to no effect. Genomically clustered Cas9 double-strand DNA breaks are associated with a strong gene-independent decrease in cell fitness. We systematically identified collateral vulnerabilities in 25% of cancer cells, introduced by tandem amplifications of tissue non-expressed genes. Our analysis demonstrates the importance of structural rearrangements in mediating the effect of CRISPR-Cas9-induced DNA damage, with implications for the use of CRISPR-Cas9 gene-editing technology, and how resulting collateral vulnerabilities are a generalisable strategy to target cancer cells.

## Introduction

Genetic loss-of-function screens are used to systematically identify genes important for cellular viability and genetic interactions in model organisms (Costanzo et al. 2010; Berns et al. 2004). Traditionally these have been performed with RNA interference (RNAi) (Marcotte et al. 2016; McDonald et al. 2017; Tsherniak et al. 2017), although its application to mammalian cells have been hampered by incomplete protein depletion and off-target effects (Jackson et al. 2006; Echeverri et al. 2006). The advent of CRISPR-Cas technologies facilitates genome engineering of human cells by addressing many of the limitations of RNAi (Shalem et al. 2014; Wu et al. 2014; Wang et al. 2014; Koike-Yusa et al. 2014) and increases capacity to identify genes essential for cellular fitness (Morgens et al. 2016; Evers et al. 2016). In cancer cell lines, CRISPR-Cas9 dropout screens have been integrated with genomic data-sets to propose novel therapeutic targets (Tzelepis et al. 2016; Marcotte et al. 2016; Hart et al. 2015; Wang et al. 2017). Tumour cells genetic instability induces synthetic-lethal dependencies on genes that otherwise have no impact on cellular fitness (Kaelin 2005). The concept of synthetic lethality has been used to selectively target cancer cells (Muller et al. 2012), translating into improvements in patient care (Farmer et al. 2005). Targeted high-throughput loss-of-function screens provide a systematic way to identify these types of cellular dependencies.

Gene copy-number changes, despite being rare and often detrimental in normal healthy cells (Itsara et al. 2009), are one of the most frequent types of genomic alterations in cancers (Beroukhim et al. 2010). They are of particular importance when analysing CRISPR-Cas9 dropout experiments, since targeting genomic regions that are copy-number amplified induces strong DNA damage responses that lead to cell cycle arrest and cell death (Aguirre et al. 2016; Munoz et al. 2016). The effect is gene-independent and ubiquitous across cancer types. This increases the false-positive rate for detecting loss of fitness (LOF) genes when interpreting CRISPR-Cas9 experiments that target amplified regions. We and others developed computational methods to account for this systematic bias (Meyers et al. 2017; Iorio et al. 2017). Some of these approaches are guided by knowledge of gene copy-number values, which on average are proportional to the non-specific LOF effect of CRISPR-Cas9 targeting. Nonetheless, the strength of this association varies significantly between cell lines and amplicons with similar copy-number, and is completely absent in some cases (Iorio et al. 2017). This indicates that other cellular features besides copy-number influence the non-specific CRISPR-Cas9 LOF effects.

Cancer cells undergo extensive genomic alterations (Sudmant et al. 2015; Li et al. 2017; Glodzik et al. 2017) but their impact on CRISPR-Cas9 targeting and response is poorly understood. Here, we combined CRISPR-Cas9 screens with whole-genome sequencing (WGS) and DNA SNP6 copy number arrays to investigate the impact of structural variation (SV) on CRISPR-Cas9 response. We find that the non-specific LOF effect of CRISPR-Cas9 screens when targeting copy-number amplified regions is specific to tandem duplications, in stark contrast to amplifications arising from chromosomal duplication. Considering cell ploidy provides more robust associations between copy-number changes and CRISPR-Cas9 LOF effects. Based on these observations, we identify cancer synthetic-lethal vulnerabilities that are elicited by targeting tandem amplified regions with CRISPR-Cas9.

## Results

### Increased cell ploidy buffers non-specific CRISPR LOF effects

We considered genome-wide CRISPR-Cas9 knockout screens performed in 17 different tumour types comprising 163 cancer cell lines which have been previously genomically characterised, i.e. copy-number and gene-expression (Supplementary Figure 1A) (Supplementary Table S1) (Behan et al. n.d.). Gene-essentiality fold-change profiles were estimated for a total of 17,234 genes, each targeted on average by 5 single-guide RNAs (sgRNAs). For the majority of the cell lines, 3 technical replicates were performed and gene log-fold change values had an average Pearson correlation (R) of 0.78. Genes previously defined as essential for cellular viability (Hart et al. 2015) were robustly recapitulated across all samples (mean Area Under Recall Curve (AURC) = 0.88), and as previously described (Munoz et al. 2016) non-detrimental genes displayed a small enrichment for positive fold-changes (mean AURC = 0.42) (Supplementary Figure 1B).

Consistent with previous findings (Aguirre et al. 2016; Munoz et al. 2016), sgRNAs targeting copy-number amplified genes were amongst those with the strongest LOF effects in the screen, even when not expressed (Figure 1A). Importantly, we observed that enrichment for LOF of sgRNA targeting amplicons varies significantly between cell lines (Figure 1B). This suggests that other factors, besides copy-number, contribute to the LOF effects found in CRISPR-Cas9. We investigated if cell ploidy differentiates cell line responses to CRISPR-Cas9 and observed that cells with higher ploidy display significantly lower fitness reduction for sgRNAs targeting copy-number amplified regions (Figure 1C). The majority of the variation observed in each copy-number group defined in (Figure 1B) can be explained by taking cell ploidy into account. Within the same cell line different chromosomes can have different number of copies, thus we estimated the number of copies of each chromosome in each cell line and assessed if this was also related with non-specific CRISPR-Cas9 LOF effects. Consistent with the ploidy status, chromosomes with more copies display significantly lower CRISPR-Cas9 LOF effects (Supplementary Figure 1C). Overall, these results show that absolute copy-number profiles need to be analysed together with cell ploidy, or chromosome copies, to model accurately the non-specific fitness reduction in CRISPR-Cas9 gene knockout experiments.

**Figure 1.**
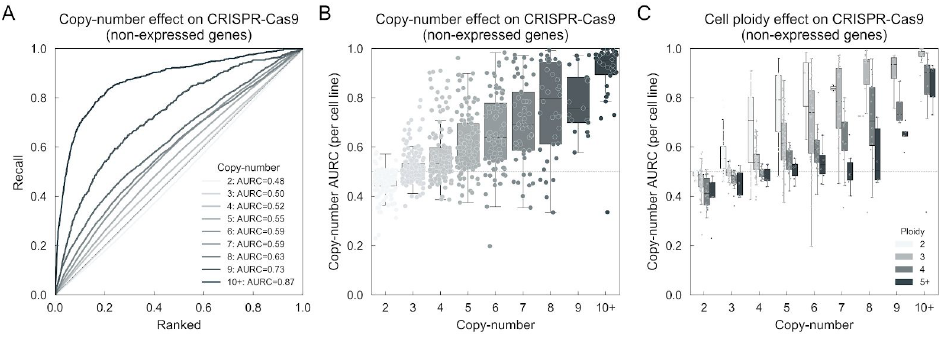
CRISPR-Cas9 screen and cell ploidy effect. (A) Enrichment of non-specific CRISPR-Cas9 LOF effects in non-expressed genes (RNA-Seq RPKM < 1) grouped by their copy-number levels, performed across all cell lines. For each copy-number group the recall curve is drawn and the AURC is reported. X-axis shows the ranked, from negative to positive, gene level CRISPR-Cas9 fold-changes. (B) Boxplots of AURCs as in A but performed in each cell line independently. Each dot represents the AURC of the given copy-number in a specific cell line. (C) Similar to B but cell lines are grouped according to their ploidy status. Boxplots represent 1.5 of the interquartile range.

### CRISPR-Cas9 LOF is specific to tandem duplications

Considering that chromosome aneuploidy is common in cancer cells, we set to analyse the implication of SV in CRISPR-Cas9 screens. To that end, we utilised WGS data of 6 breast cancer cell lines with a matched normal to call somatic SVs, such as tandem duplications, translocations, deletions and inversions, using BRASS (BReakpoint AnalySiS) (Glodzik et al. 2017). Tandem duplications were the most frequent type of rearrangements across the 6 cell lines (Figure 2A), recapitulating previous observations that this is a frequent event in breast cancers (McBride et al. 2012; Glodzik et al. 2017). Utilising a statistical framework, we then searched for associations between SVs and CRISPR-Cas9 LOF effects. SVs were most informative of CRISPR-Cas9 response when accompanied by copy-number alterations, with strong LOF falling within tandem duplications (Figure 2 B, C and D). Nested tandem duplications (Figure 2B and Supplementary Figure 2A) and complex patterns of SVs with large number of translocations (Supplementary Figure 2 B, C) were also visible and these comprised some of the strongest LOF responses. Importantly, copy-number amplifications not overlapping with tandem-duplications were in general not associated with LOF effects (Supplementary Figure 2D). These examples show how SVs, specifically tandem duplications, result in copy-number alterations that lead to non-specific LOF effects in CRISPR-Cas9 experiments.

**Figure 2.**
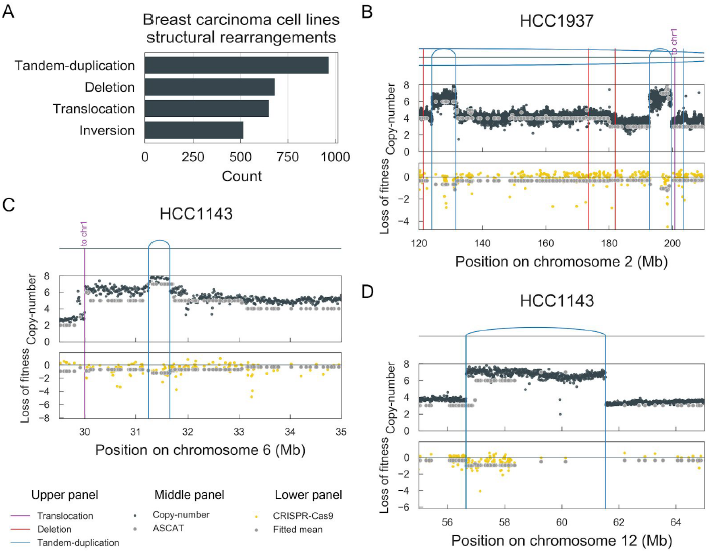
Structural variation impacts CRISPR-Cas9 response. (A) Count of somatic structural rearrangements identified in 6 breast cancer cell lines. (B), (C) and (D) are representative examples of the strongest associations between SVs and CRISPR-Cas9 LOF. Structural rearrangements are mapped in the upper panel, in the middle panel copy-number levels are represented and in the lower panel CRISPR-Cas9 gene level fold-changes are shown. SVs are coloured with tandem duplications defined with blue lines, deletions with red lines and chromosome translocations in purple. Average mean values for copy-number (middle panel) and responses to CRISPR-Cas9 (lower panel) are represented as gray circles.

To more comprehensively investigate the impact of tandem duplications in CRISPR-Cas9 experiments, we propose to normalise gene copy-numbers by the chromosome copy-numbers, termed hereafter as gene copy-number ratio, on an individual cell line basis (Figure 3A, Supplementary Table 2). This was necessary because WGS was not available for most cell lines. The ratio encompasses three scenarios: a value (i) less than 1 represents a gene depletion, (ii) equal to 1 represents either a normal diploid chromosome with 2 copies of the gene or deletions/amplifications that are consistent in the gene and the chromosome; (iii) greater than 1 represents genes that have been amplified more than the chromosome to which they map, likely representing tandem or interspersed duplications. This ratio allows us to separate gene duplications that originate from whole chromosome/genome duplication from those arising from smaller tandem amplifications, which we hypothesize induces stronger CRISPR-Cas9 LOF effects.

**Figure 3.**
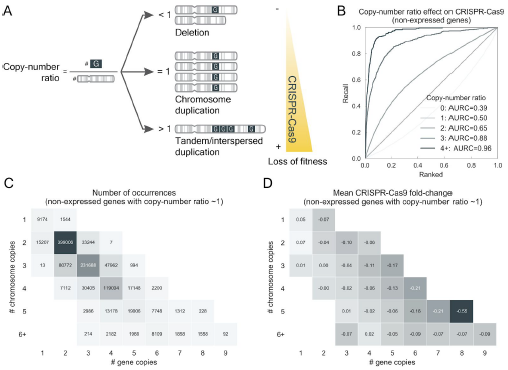
Gene copy-number ratio association with CRISPR-Cas9 loss of fitness effect. (A) Diagram of the different genomic rearrangements captured by the gene copy-number ratio and their potential effect in CRISPR-Cas9 response. (B) Enrichment of high copy-number ratios to strong CRISPR-Cas9 LOF effects. Recall curves of non-expressed genes grouped by their copy-number ratio profile across CRISPR-Cas9 fold-changes. (C) Matrix of gene and chromosome copies. The total number of events in each condition with a copy number ratio equal to 1 (rounded to nearest whole number) are shown, and (D) represents the mean gene-level CRISPR-Cas9 fold-change (log2) in the respective group.

Non-expressed genes with copy-number ratios greater than 1 showed strikingly higher LOF effects (Figure 3B, Supplementary Figure 3A). Ratios greater than 4 displayed among the strongest LOF effects captured in the screen (mean log2 fold-change <-1) (Supplementary Figure 3A). The distribution of the copy-number ratios across all cell lines was centered around 1 confirming that the vast majority of copy-number alterations originate from whole chromosome duplications (Supplementary Figure 3B). Notably, we observed thousands of occurrences of highly amplified genes with a copy-number ratio close to 1 which displayed almost no CRISPR-Cas9 LOF effects (Figure 3 C, D). Copy-number ratios displayed no association with LOF responses obtained with RNAi experiments (McDonald et al. 2017) (Supplementary Figure 3C), confirming this is a feature specific of CRISPR-Cas9 screens. Performing Fluorescence *In Situ* Hybridization (FISH) in two *MYC* amplified cell lines with distinct copy-number ratios, we confirmed that high copy-number ratios represent strong focal tandem amplifications (Supplementary Figure 3 D, F), with more defined CRISPR-Cas9 LOF effects (Supplementary Figure 3 E, G). Systematic analysis of copy-number ratios relies on accurate cell ploidy and chromosome copy-number estimations from WGS or SNP6 arrays, thereby it is notable that copy number ratio estimates were highly consistent with FISH karyotypes (Supplementary Figure 4 A, B).

Overall, these results are important for the analysis of CRISPR-Cas9 datasets showing that non-specific LOF effects induced by targeting of copy-number amplified regions are specific to tandem duplicated regions, while copy-number amplifications originating from chromosome duplication have little to no effect.

### CRISPR-Cas9 collateral vulnerabilities in cancer cells

Our finding that targeting tandem amplifications with CRISPR-Cas9 induces a potent and specific LOF effect could facilitate exploitation of this mechanism to develop targeted cancer therapies. Cancer cells generally display profound alterations of their karyotype with complex genomic rearrangements, which are rare in healthy cells. Targeting copy number amplified regions with CRISPR-Cas9 to induce synthetic-lethal effects has been postulated (Aguirre et al. 2016). Hence, here we systematically investigate the prevalence of collateral vulnerabilities and refine this concept to targeting tandem duplicated genomic regions with CRISPR-Cas9 reagents, as these are likely to have strong and selective LOF effects in cancer cells. We focus on tandem duplicated regions, which likely but not necessarily contain oncogenes, and non-expressed genes to limit potential toxicity in healthy tissue (Figure 4A). We refer to this concept as CRISPR-Cas9 collateral essentiality.

**Figure 4.**
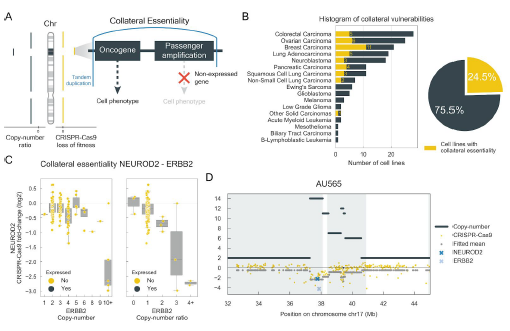
CRISPR-Cas9 collateral essentialities. (A) Schematic representation of collateral essentialities identified from tandem duplicated regions (high copy-number ratio) which likely, but not necessarily, contain oncogenes and genes that are passenger amplified. CRISPR-Cas9 targeting of passenger amplified genes should not be detrimental for normal cells if they are not expressed. (B) Distribution per cancer type of significant collateral essentialities. Cell lines highlighted (yellow) contain at least one collateral essentiality. (C) Most frequent collateral essentiality found in the screen. Left panel relates gene copy-number profile while right panel relates gene copy-number ratio of ERBB2 to NEUROD2 CRISPR-Cas9 fold-change. (D) Representation of both copy-number and LOF of the collateral essentiality in the cell line AU565. Copy-number represent absolute copy-number estimated and CRISPR-Cas9 represents gene level LOF (log2) fold-changes.

We systematically searched the 163 cancer cell lines for collateral essentialities and used stringent filters to focus on genes that: (i) have strong LOF effects; (ii) are genomically colocalized with a high copy-number ratio gene; and (iii) are not expressed across the majority of the cell lines of the same tissue. A total of 223 CRISPR-Cas9 collateral essentialities were identified (Supplementary Table 3) (194 unique genes), 59.2% of these involved associations with 53 cancer driver genes identified in COSMIC (Forbes et al. 2015). Collateral essentialities were identified in 24.5% of the screened cell lines (40 cell lines across 9 different tumour types), demonstrating the general nature of this mechanism (Figure 4B).

Among the most frequent collateral essentialities was *NEUROD2* associated with *ERBB2* amplifications present in 4 cell lines (Figure 4C), and *MRGPRD* associated with *CCND1* amplifications present in 3 cell lines (Supplementary Figure 5A, B). *NEUROD2* is involved in neuronal differentiation, mainly expressed in the brain (Supplementary Figure 5C), and frequently amplified in breast tumours (Supplementary Figure 5D). Focusing on the cell line with the strongest amplification of *ERBB2* and strongest LOF response to *NEUROD2* knockout, i.e. breast carcinoma cell line *AU565*, we confirmed that *ERBB2* and *NEUROD2* are co-genomically amplified (Figure 4D). Notably, gene copy-number ratio separates better the CRISPR-Cas9 LOF responses compared to the absolute copy-number (Figure 4C), suggesting that some cell lines do not respond to *NEUROD2* knockout likely due to *ERBB2* amplifications arising from chromosome duplications. Together, these results illustrate the concept of selectively targeting cancer cells with potentially reduced toxicity by exploiting CRISPR-Cas9 collateral essentiality.

### *Crispy*, a flexible tool to identify genomic determinants of CRISPR-Cas9 response

Lastly, for this study we developed an open-source Python module termed *Crispy* to systematically identify associations between genomic alterations, e.g. SVs, and CRISPR-Cas9 screens (Supplementary Figure 6A). Contrary to methods that need to be trained across panels of different cell lines, *Crispy* is trained on a per chromosome and per sample basis. This way the variability in CRISPR-Cas9 LOF effects due to chromosome aneuploidies are modeled independently for each chromosome providing more accurate estimations (Supplementary Figure 6 B, C), while preserving the capacity to identify LOF genes (Supplementary Figure 6D). Of note, *Crispy* takes into consideration that high copy-number amplifications might have no impact if arising from whole chromosome amplifications, avoiding potential over-correction of CRISPR-Cas9 fold-changes. *Crispy* quantifies the impact of genomic alterations in CRISPR-Cas9 response and uses this to calculate CRISPR-Cas9 fold-changes that are corrected for potential biases, including copy-number amplifications in tandem duplicated regions.

## Discussion

In this study, we demonstrate that copy-number amplifications lead to non-specific CRISPR-Cas9 LOF effects if originating from tandem duplications. In contrast, no impact is observed from gene amplifications associated with increased cell ploidy. We devised a gene copy-number metric, normalised by chromosome copy-number, that greatly improves the ability to classify which copy-number amplifications will result in a LOF bias in CRISPR-Cas9 experiments. Notably, our findings were recapitulated when considering only non-expressed genes, emphasising that the LOF bias is not due to a potential biological function of the genes. Combining the copy-number ratios, WGS and FISH experiments, we validated that tandem duplications are the most frequent SVs associated with CRISPR-Cas9 non-specific deleterious effects. Nonetheless, we cannot exclude that other events might also play a role. In particular, extrachromosomal DNAs (ecDNA) have been found widespread in cancer (Stephens et al. 2011; Turner et al. 2017) and are tandem duplicated rich DNA sequences, although we did not observe any evidence of ecDNAs in the cell lines analysed with FISH.

We hypothesise that CRISPR-Cas9 non-specific LOF effects in copy-number amplified regions is buffered if Cas9 double-strand breaks (DSB) are spread across several chromosomes (Supplementary Figure 7). We expect that nonhomologous end joining (NHEJ) repair of clustered DSBs can lead to large deletions and changes in copy-number, thereby inducing strong DNA damage and LOF responses. Hence, chromosomal structural alterations such as tandem duplications can determine CRISPR-Cas9 response. Moreover, this presents evidence that structural rearrangements can impact the functioning of DNA damage repair mechanisms with important implications in cellular phenotype.

Tandem duplications are among the most frequent structural rearrangements in cancer (Li et al. 2017) indicating that our findings are of general importance when designing and interpreting CRISPR-Cas9 experiments. Specifically, targeting genes that reside within regions of tandem duplication, whether knocking out individual genes, performing genetic screens using a library of sgRNAs, or performing specific gene edits, will lead to strong non-specific LOF effects. Computational methods such as *Crispy* and others (Iorio et al. 2017) can efficiently correct for this bias in CRISPR-Cas9 screens. However, this is unlikely to be possible for many studies, for example the common scenario of a single gene knockout or alteration in individual cell lines. In these cases, information about gene copy number, cell line ploidy and ideally SV information should be incorporated. We expect that the bias in CRISPR-Cas9 data described here is a general phenomenon and consequently will be observed in other cancer cell models such as patient-derived xenografts and organoids, and potentially also present in other types of CRISPR-Cas based systems that introduce DNA double-strand breaks.

Collateral vulnerabilities are typically defined as cancer-specific therapeutic vulnerabilities offered by passenger gene deletion or inactivation of non-tumour suppressor genes, and this is a well-established paradigm in oncology. Here we extend this concept to cancer-specific vulnerabilities that can be exploited through CRISPR-Cas9 targeting of tandem duplicated regions of cancer genomes, and demonstrate that this a general phenomenon in cancer cells. This theoretical approach could be applied based on the detailed analysis of an individual genome, or by targeting regions of recurrent tandem duplication (Glodzik et al. 2017). We have examined the application to non-expressed genes, but this may be effective when targeting non-genic regions, further reducing the risk of toxicity to healthy cells. CRISPR-Cas9 applications to gene editing and personalised medicine are in their infancy but quickly progressing, and promising results have been obtained by injecting localised viral vectors expressing CRISPR-Cas (Xue et al. 2016). Nonetheless, important hurdles remain with respect to delivery, efficiency and safety before this approach could be considered clinically. In summary, this work provides new insights into CRISPR-Cas9 mediated LOF effects, with implication for the use of CRISPR-Cas9 gene-editing, and points to CRISPR-Cas9 collateral essentiality as an approach to develop tumour cell specific therapies.

## Acknowledgements

We gratefully acknowledge Pedro Beltrao, Dominik Glodzik, Helen Davies and Simon Brunner for helpful comments and discussions. Work in M.G lab is supported by funding from Open Targets (OTAR015), CRUK (C44943/A22536), SU2C (SU2C-AACR-DT1213) and the Wellcome Trust (102696).

## Author contributions

Conceptualization: E.G., F.I., K.Y., M.G.

Software: E.G., D.A.

Validation: E.G., D.A., F.I., M.G.

Formal analysis: E.G., D.A.

Investigation: F.B., S.L., F.Y.

Resources: M.G.

Data curation: E.G., F.I.

Writing - original draft preparation: E.G., M.G.

Writing - reviewing and editing: E.G., M.G., K.Y., F.I., F.B., F.Y., D.A., E.S.

Visualisation: E.G.

Supervision: F.I., K.Y., E.S., O.S., M.G.

Funding acquisition: K.Y., M.G.

## Materials and Methods

### Processing of CRISPR-Cas9, SNP6 and RNA-seq samples

A CRISPR-Cas9 library containing 95,994 sgRNAs was utilised (Koike-Yusa et al. 2014; Tzelepis et al. 2016) to screen the loss of fitness (LOF) impact of knocking-out each 17,234 genes across 163 cell lines as described here (Behan et al. n.d.). Raw sequence counts of each sgRNA were corrected by library size in each sample using median-of-ratios method, similar to DESeq2 (Anders & Huber 2010). Only replicates with a Pearson correlation (R) coefficient greater than 0.7 were considered. Non-targeting plasmid control sample was used and sgRNAs with lower than 30 counts were discarded. Log2 sgRNA fold-changes were estimated between samples and the plasmid control. Gene level estimates of the fold-changes were calculated by averaging all mapping sgRNA fold-changes. Single nucleotide polymorphism (SNP) array hybridization using the Affymetrix SNP6.0 platform was performed according to Affymetrix protocols. Segment copy-number variants were obtained using PICNIC (Greenman et al. 2010) as previously described (Iorio et al. 2016). RNA-seq experiments for CRISPR-Cas9 profiled cell lines were assembled from multiple data-sets (Garcia-Alonso et al. 2017). To minimise technical bias, all samples were processed with the same pipeline, iRAP (Fonseca et al. 2014), to obtain raw counts. Genes with Reads Per Kilobase per Million (RPKM) with zero counts were termed as non-expressed in the particular sample.

### Chromosome harvest and fluorescence *in situ* hybridisation (FISH)

Metaphase chromosomes were harvested from the cancer cell lines after incubation with 0.05 g/ml of colcemide (Thermo-Fisher) for 2-3 h. Subsequently, cells were treated with a buffered hypotonic solution (0.4% KCl in 10 mM HEPES, pH7.4) for 8 - 12 min at 37°C and fixed with 4:1(v/v) methanol:glacial fixative. The human fosmid clone WI2-1694H13 was labelled with green-dUTP as described in (Stephens et al. 2011). Human 24 colour FISH (M-FISH) probe preparation and slides treatments followed (Agu et al. 2015) with slight modifications. Freshly-prepared metaphase slides were immersed in acetone for 10 min and then baked at 62 C for 1 hour. Slides were denatured in an alkaline denaturation solution (0.5 M NaOH, 1.5 M NaCl, Sigma-Aldrich) for 9-10 min. Metaphases were examined with a Zeiss AxioIamger D1 fluorescence microscope. FISH images were captured using the SmartCapture software (Digital Scientific UK) and karyotyped using the SmartType Karyotyper software (Digital Scientific UK). Ten metaphases for each sample were analysed by M-FISH.

### Whole-genome sequencing

DNA of 6 cancer cell lines and 6 EBV derived matched normal cell lines were obtained and sequenced with massively parallel Illumina sequencing technology (EGAD00001004124) and aligned to the human reference genome GRCh37 using Burrows-Wheeler Aligner (v0.5.9) (Li & Durbin 2009). Average sequence coverage was 43-fold for cancer cell lines and 42-fold for matched normal. Somatic structural rearrangements were identified by providing aligned bam files to BRASS (BReakpoint AnalySiS) (https://github.com/cancerit/BRASS/). BRASS calls structural variations via assembly of discordant paired-end reads. ASCAT (v4.0.1) (Van Loo et al. 2010) was used to perform copy-number variation analysis. For fitting with Crispy, SVs were discretized, each gene covered by the CRISPR-Cas9 screen was marked with a 1 if its genomic region overlapped with a tandem duplication, inversion or deletion, 0 otherwise, generating a binary table of 3 columns stating the SVs mapping to each gene.

### *Crispy* Python module to identify genomic determinants of CRISPR-Cas9 screens

*Crispy* is a flexible tool to identify systematically associations between multiple types of genomic alterations and CRISPR-Cas9 response. Gaussian processes regressions implemented on scikit-learn Python module (v0.19.1) (Pedregosa et al. 2011) were used, specifically a squared-exponential kernel with a length scale (θ) hyperparameter varying between 1e-5 and 10 is used. A constant (σ) and noise (ψ) kernels are also added:

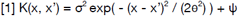

Where θ determines the length of waves and the σ defines the average distance from the mean. Fitting is performed at the gene level, and the input to the kernels are: (i) an integer array with the copy-number profiles, (ii) concatenated with a binary matrix identifying the SVs that overlap with the gene genomic region. The genomic features are used to model the CRISPR-Cas9 gene log fold-changes. Default configurations of scikit-learn Gaussian regression are used except n_restarts_optimizer is set to 3, to initialise the optimisation procedure multiple times. The complex kernel defined in [1] is fitted independently in each chromosome and each sample. Specifically, genes and genomic alterations are grouped into the different chromosomes they map to. This guarantees that CRISPR-Cas9 global effects that are sample or chromosome specific are captured automatically by the defined kernel [1]. *Crispy* takes advantage of this modular approach and allows each chromosome to be processed in parallel (parallel computing) using native Python parallelizing packages, reducing considerably the execution time. After optimising the kernel [1] CRISPR-Cas9 corrected fold-changes are obtained by subtracting from the original fold-changes the predicted bias from the inputted genomic alterations. We systematically tested *Crispy* across all 163 cell lines providing as input the gene copy-number profiles. Reassuringly, *Crispy* corrected fold-changes successfully corrected the majority of the copy-number induced LOF CRISPR-Cas9 bias (Supplementary Figure 6B) and preserved the capacity to identify essential genes (Supplementary Figure 6D). Moreover, the ploidy association with LOF responses was greatly reduced after *Crispy* correction (Supplementary Figure 6C).

### Gene copy-number ratios

Gene copy-number ratios, i.e. number of absolute gene copies divided by the number of copies of the respective chromosome, are performed for all protein-coding genes annotated in the human GRCh37 reference assembly. Segment absolute copy-numbers are mapped to the assembly using BEDtools (v2.27.1) (Quinlan & Hall 2010) and pybedtools (v0.7.10) (Dale et al. 2011). Gene and chromosome absolute copy-number values are estimated by taking the copy-number weighted mean of all the mapping segments weighted by their size. Gene copy-number ratios across the 163 cell lines are available in Supplementary Table 2.

To verify that high copy-number ratios represent strong focal chromosome amplifications we applied FISH and probed the location of the frequently and highly amplified oncogene *MYC*. We chose 2 cell lines (HCC1954 and NCI-H2087) with high *MYC* absolute copy-number (9 and 7, respectively) but discordant copy-number ratios (1.58 and 4.05, respectively) due to different ploidy. Consistent with our hypothesis, *MYC* amplifications in the tetraploid cell line (HCC1954) displays a small level of tandem duplications of *MYC* while the majority of the copies were spread across derivatives of chromosome 8 (Supplementary Figure 3D). In contrast, in the diploid cell line (NCI-H2087) *MYC* copies were mostly concentrated in one arm of a derivative of chromosome 8 (Supplementary Figure 3F). A control triploid cell line (LS1034) with diploid *MYC* and rounded copy-number ratio 1 was analysed and corroborated our prediction that chromosome 8 is mostly diploid and contains 2 copies of *MYC* (Supplementary Figure 4A).

### Collateral essentialities

Collateral essentialities were identified across all 163 cancer cell lines using CRISPR-Cas9 log2 fold-changes, SNP6 estimated copy-number ratios and RNA-Seq defined non-expressed genes. Firstly, tandem amplified genes across cell lines were defined as those with a copy-number ratio greater than 2. Secondly, non-expressed essential genes were defined in each cell line as those genes with a RPKM < 1 and a CRISPR-Cas9 log fold-change lower than the mean of known essential genes (Hart et al. 2015). An off-set of 25% of the mean log fold-change of essential genes was used in order to capture genes with strong LOF effects that are not as strong as essential genes. In order to exclude genes with a strong LOF across most of the cell lines and focus on those that are tissue or subtype specific, we excluded genes that had stronger LOF effects than the offsetted essential genes in more than 15% of the total cell lines (24 cell lines). Lastly, collateral essentialities were defined as pairs of genes that in the same cell line one is tandem amplified and the other is non-expressed and essential. Only gene pairs that are on the same chromosome and genomically co-localised were considered. The genomic distance between the two genes has to be lower than 1 Mb. Vulnerable cell lines were defined as those with at least one collateral essentiality and for which the collateral essential gene (non-expressed and essential) was not expressed in at least 50% of the cell lines from the same tissue, thus reducing tissue specific essential genes and potential cytotoxic effects. Copy-number amplification data for the relevant genes in tumour samples was obtained from cBioPortal (Cerami et al. 2012; Gao et al. 2013) (accession date 31/03/2018) filtering for BRCA cohorts and only copy-number alterations. Gene expression measurements in healthy tissues was obtained from the GTex data-base (GTEx Consortium 2013), specifically gene median transcript per million (TPM) per tissue.

### Code availability

*Crispy* is a Python module (https://github.com/EmanuelGoncalves/crispy) and its code is distributed under the open-source 3-Clause BSD License.

## Supplementary Figures

**Supplementary Figure 1.**
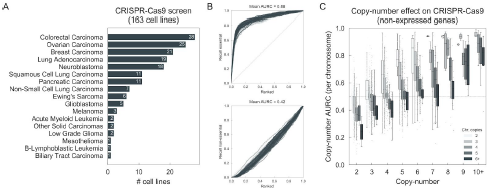
Screen, quality control and loss of fitness bias of CRISPR-Cas9 fold-changes. (A) Number of cell lines screened per cancer type. (B) Recall curves of previously defined core-essential (top) and non-essential (bottom) genes (Hart et al. 2015). X-axis represents the ranked CRISPR-Cas9 gene-level fold-changes from negative to positive. AURC of each curve was estimated and the mean is reported. C) AURC of non-expressed genes (RNA-seq RPKM < 1) estimated per chromosome in each cell line independently. Chromosomes were grouped according to their estimated number of copies.

**Supplementary Figure 2.**
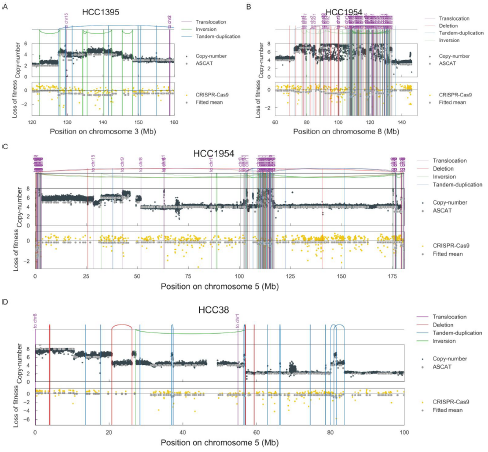
SV associations with CRISPR-Cas9 response. Representative cases of the strongest associations between SVs and CRISPR-Cas9 loss of fitness involving (A) nested tandem duplications; (B) and (C) complex patterns of SVs containing a high number of translocations and aligned with the copy-number changes; and (D) lack of non-specific loss of fitness effects in copy-number amplified regions that are not associated with tandem duplications.

**Supplementary Figure 3.**
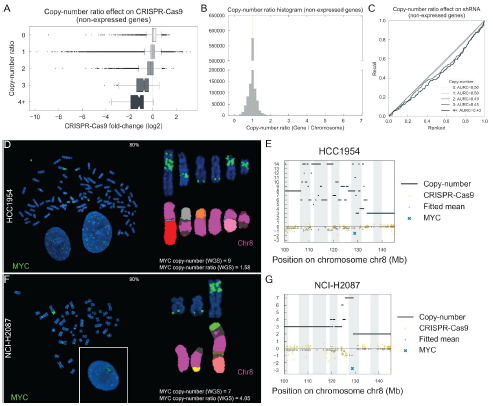
Gene copy-number ratios. (A) CRISPR-Cas9 fold-changes of non-expressed genes grouped according to their copy-number ratio. (B) Gene copy-number ratios distribution across all cell lines. (C) Recall curves of shRNA LOF scores of non-expressed genes grouped by their copy-number ratio profiles. (D) FISH of MYC amplifications (green signal) in HCC1954 tetraploid cell line. In the left panel, representative metaphase (80% of cells) with high MYC amplifications and low copy-number ratio. In the right panel, detailed view of the chromosomes containing MYC signal (upper) and the corresponding derivative chromosomes identified by M-FISH (lower). (E) Genomic region containing MYC and the copy-number profile (SNP6) and CRISPR-Cas9 log fold-changes in HCC1954. (F) FISH of MYC amplifications (green signal) in NCI-H2087 diploid cell line. In the left panel, representative metaphase (90% of cells) with high MYC copy-number ratio. In the right panel, detailed view of the chromosomes containing MYC signal (upper) and the corresponding derivative chromosomes identified by M-FISH (lower). (G) Similar to E, genomic region containing MYC displaying copy-number and CRISPR-Cas9 measurements. For both cell lines 10 cells were analyzed in the FISH experiments.

**Supplementary Figure 4.**
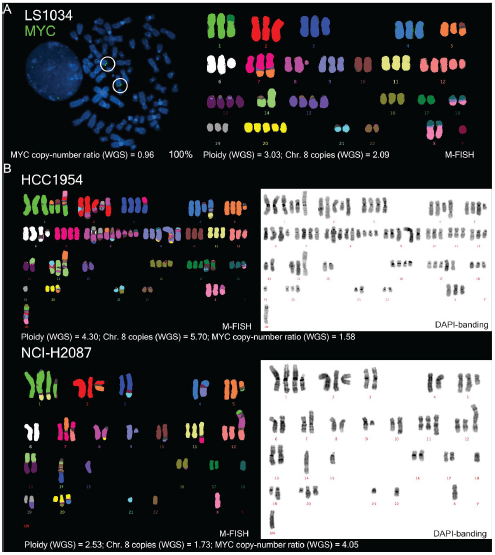
FISH and M-FISH experiments. (A) FISH with green fluorescent MYC probe in triploid cell line with 2 MYC copies (left). M-FISH representative karyotype (right). 10 cells were used and the karyotype was observed in 100% of the cells. (B) M-FISH representative karyotypes (10 cells) of two MYC amplified cell lines. MYC copy-number ratio, ploidy and chromosome copy-number were estimated from WGS.

**Supplementary Figure 5.**
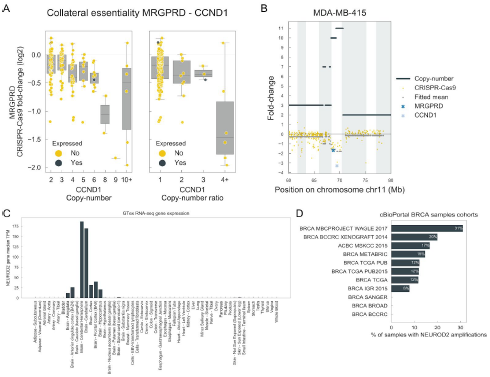
Collateral essentialities. (A) Representative example of one of the most frequent collateral essentialities found and (B) displays the genomic region containing the associated genes. (C) NEUROD2 expression in healthy cells. (D) frequency of NEUROD2 copy-number amplification in breast tumour samples.

**Supplementary Figure 6.**
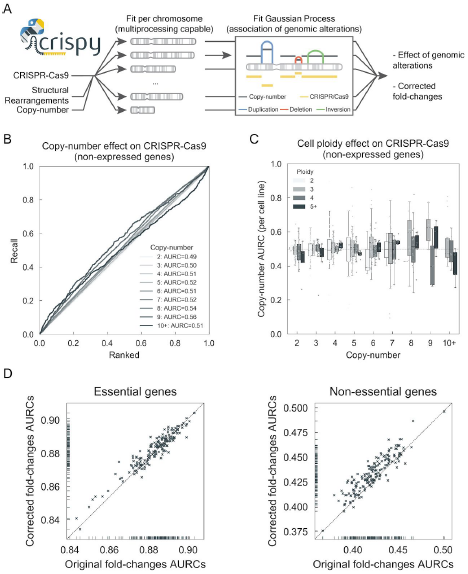
Crispy method and benchmark. (A) Diagram of the workflow implemented in Crispy integrating different types of genomic alterations and CRISPR-Cas9 response. (B) Recall curves of non-expressed genes (RNA-seq RPKM < 1) using Crispy corrected CRISPR-Cas9 fold-changes. Genes were grouped by their copy-number profile. C) Boxplots of AURC performed in each cell line independently using Crispy corrected fold changes. Cell lines were grouped by their ploidy status. D) Comparison between AURCs of previously defined (Hart et al. 2015) core-essential (left) and non-essential (right) obtained using CRISPR-Cas9 fold-changes for original and Crispy corrected CRISPR-Cas9 fold-changes.

**Supplementary Figure 7.**
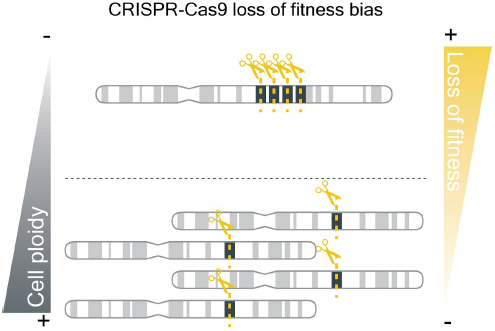
Graphical abstract representing different types of genomic amplifications and their impact on CRISPR-Cas9 loss of fitness (LOF) responses. CRISPR-Cas9 LOF effects induced by copy-number amplifications is buffered by increased cell ploidy and chromosome copy-number due to spread of the genomic cuts. Tandem duplications lead to high density of CRISPR-Cas9 double strand breaks (DSB) in the same genomic region which is associated with stronger non-specific LOF responses, in contrast to similar numbers of DSBs but arising across multiple chromosomes.

## Supplementary Table legends

**Supplementary Table 1.** List of cancer cell lines included in the study indicating the tissue of origin, experimental data available and ploidy status derived from SNP6 arrays.

**Supplementary Table 2.** Genome-wide copy-number ratios estimated from SNP6 arrays for all cell lines.

**Supplementary Table 3.** List of collateral essentialities found across the cell lines.

